# Assembling Large Genomes with Single-Molecule Sequencing and Locality Sensitive Hashing

**DOI:** 10.1101/008003

**Authors:** Konstantin Berlin, Sergey Koren, Chen-Shan Chin, James Drake, Jane M. Landolin, Adam M. Phillippy

**Affiliations:** Department of Chemistry and Biochemistry, University of Maryland, College Park, MD 20742, USA; Institute for Advanced Computer Studies, University of Maryland, College Park, MD 20742, USA; National Biodefense Analysis and Countermeasures Center, Frederick, MD 21702, USA; Pacific Biosciences, Menlo Park, CA 94025, USA

## Abstract

We report reference-grade *de novo* assemblies of four model organisms and the human genome from single-molecule, real-time (SMRT) sequencing. Long-read SMRT sequencing is routinely used to finish microbial genomes, but the available assembly methods have not scaled well to larger genomes. Here we introduce the MinHash Alignment Process (MHAP) for efficient overlapping of noisy, long reads using probabilistic, locality-sensitive hashing. Together with Celera Assembler, MHAP was used to reconstruct the genomes of *Escherichia coli*, *Saccharomyces cerevisiae*, *Arabidopsis thaliana*, *Drosophila melanogaster*, and human from high-coverage SMRT sequencing. The resulting assemblies include fully resolved chromosome arms and close persistent gaps in these important reference genomes, including heterochromatic and telomeric transition sequences. For *D. melanogaster*, MHAP achieved a 600-fold speedup relative to prior methods and a cloud computing cost of a few hundred dollars. These results demonstrate that single-molecule sequencing alone can produce near-complete eukaryotic genomes at modest cost.

## Introduction

Genome assembly is the process of reconstructing a genome from a collection of much shorter sequencing reads, and is a critical step of all genome projects^1–3^. In contrast to resequencing projects, *de novo* assembly is not assisted by a reference and the genome must be reconstructed from scratch. An accurate reconstruction is critical, as both the continuity and base accuracy of an assembly can affect the results of all downstream analyses. However, repetitive sequences make assembly a difficult problem when the repeat length exceeds the read length^4, 5^. Unfortunately, most high-throughput sequencing methods generate sequencing reads of only a few hundred base pairs, which is well short of many common repeats. While short reads are sufficient for many types of analyses, they are not sufficient for assembling through the major repeat families in both microbial and eukaryotic genomes. This has led to more fractured and incomplete assemblies^6^ compared to those based on first-generation Sanger sequencing^7^.

Recent advances in single-molecule sequencing technologies have promised reads hundreds of fold longer than second-generation methods^8–11^. Most notably, Pacific Biosciences’ single-molecule, real-time (SMRT^®^) sequencing was the first commercially available long-read technology^11^. Utilizing a DNA polymerase anchored in a zero-mode waveguide, SMRT sequencing has delivered usable reads upwards of 50Kbp^12^. Preliminary results from Oxford Nanopore suggest that >10Kbp read lengths are also possible using nanopore sequencing^13, 14^. Such read lengths drastically simplify genome assembly by resolving repetitive structures in the assembly graph^12, 15^. However, the long reads generated by single-molecule sequencing currently suffer from low accuracy, requiring new algorithms to compensate for noise in individual reads. Recent theoretical work has demonstrated that random error can be overcome algorithmically^16^, and early studies indicate that SMRT sequencing follows a largely random error model^17, 18^. Thus, by oversampling the genome at sufficient coverage, SMRT sequencing can be used to produce assemblies of unmatched continuity^15, 17, 19–21^, including automatically finished genomes for the majority of known bacteria and archaea^15^.

Despite providing outstanding results, early assemblies of noisy, long reads have come at a substantial computational cost. For example, using tools available at the time, an initial assembly of *Drosophila melanogaster* from SMRT reads required over 600,000 CPU hours—the equivalent of more than 20 days running on a thousand-core compute cluster^22^. Even small bacterial genomes currently require a day to assemble using the HGAP^19^ or PBcR^15^ assembly pipelines. Until now, the primary bottleneck of long-read assembly has been the sensitive all-versus-all alignment required to determine overlapping read pairs. For the initial *D. melanogaster* assembly, this overlapping step comprised over 95% of the total runtime. Even if future long-read technologies improve accuracy, all-pairs overlapping would remain a significant bottleneck in overlap-layout-consensus (OLC) assembly^1^. Both this computational cost and the comparatively high sequencing cost have prevented widespread application of SMRT sequencing to large genomes. The steadily increasing throughput of the PacBio^®^ RS II instrument has begun to address sequencing costs, but the computational cost of assembling larger genomes has remained beyond the reach of most investigators.

To address the computational barrier of long-read assembly, we present a probabilistic algorithm for efficiently detecting overlaps between noisy, long reads. The MinHash Alignment Process (MHAP) uses a dimensionality reduction technique called MinHash^23^ to create a more compact representation of sequencing reads. Originally developed to determine the similarity of web pages^24^, MinHash reduces a text or string to a small set of fingerprints, called a sketch. MinHash sketches have been successfully applied to document similarity^23^, image similarity^25^, sequence similarity^26, 27^, and metagenomic clustering^28^. The approach can also be viewed as a generalization of minimizers^29^. Briefly, to create a sketch for a DNA sequence, all *k*-mers (aka shingles or *q*-grams) are converted to integer fingerprints using multiple, randomized hash functions. For each hash function, only the minimum valued fingerprint, or min-mer, is retained. The collection of min-mers for a sequence makes the sketch (Fig. 1, Methods). This locality sensitive hashing allows the Jaccard^30^ similarity of two *k*-mer sets to be estimated by simply computing the Hamming^31^ distance between their sketches. Because the sketches are small and Hamming distance is quick to compute, this is an extremely efficient technique to estimate similarity.

**Figure 1.**
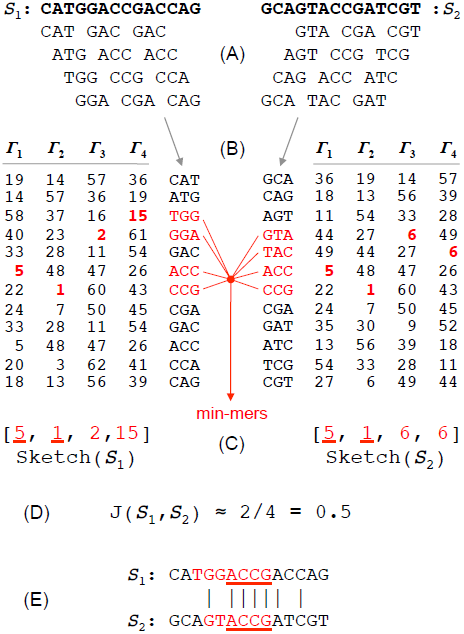
**Rapid overlapping of noisy reads using MinHash sketches** To create a MinHash sketch of a DNA sequence *S*, (A) the sequence is first decomposed into its constituent *k*-mers. In this toy example, *k*=3, resulting in 12 *k*-mers each for *S*_1_ and *S*_2_. (B) All *k*-mers are then converted to integer fingerprints via multiple hash functions. The number of hash functions determines the resulting sketch size *H*. Here *H*=4. *H* independent hash sets are generated for each sequence (*Γ*_1..*H*_). In MHAP, after the initial hash (*Γ*_1_), subsequent fingerprints are generated using an XORShift pseudorandom number generator (*Γ*_2..H_). The *k*-mer generating the minimum value for each hash is referred to as the min-mer for that hash. (C) The sketch of a sequence is composed of the ordered set of its *H* min-mer fingerprints. In this example, the sketches of *S*_1_ and *S*_2_ share the same minimum fingerprints for *Γ*_1_ and *Γ*_2_. (D) The fraction of entries shared between the sketches of two sequences *S*_1_ and *S*_2_ (0.5) serves as an estimate of their true Jaccard similarity (0.22), with the error bound controlled by *H*. In practice, *H* >> 4 is required to obtain accurate estimates. (E) If sufficient similarity is detected between two sketches, the shared min-mers (ACC and CCG in this case) are located in the original sequences and the median difference in their positions is computed to determine the overlap offset (0) for *S*_1_ and *S*_2_.

Here, we present assemblies of SMRT sequencing reads for four model organisms and the human genome using a MHAP-enabled assembly pipeline. The resulting assemblies are superior to any previous *de novo* assemblies of these organisms, and were produced in a fraction of the time required using previous overlapping algorithms^32^. In addition, these assemblies include novel heterochromatic sequences and fill persistent gaps remaining in the reference genomes of these important organisms.

## Results

### MinHash alignment filtering

MHAP uses MinHash sketches for efficient alignment filtering. The time required to hash, index, store, and compare *k*-mers is proportional to the sketch size, so it is preferable to keep sketches small. However, using fewer min-mers reduces the sensitivity of the filter. Figures 2a and 2b show that it is possible to use sketches an order of magnitude smaller than the input reads, while maintaining acceptable overlap detection accuracy. Specifically, sketches of 1,000 16-mers can be used to accurately detect 5Kbp overlaps for 10Kbp reads simulated from the human genome with an overlap error rate of 30% (Methods, Supplementary Note 1). For human, using a smaller value of *k* (e.g. 10) increases the number of false matches found, so it is preferable to use the largest value of *k* that maintains sensitivity. Sensitivity can be further controlled by the sketch size. Because the error rate of an alignment is roughly additive in the error rate of the two reads, mapping high-error reads to a reference genome is easier than overlapping. For mapping 10Kbp reads to the human genome at 15% error, a sketch of only ∼150 16-mers is required to achieve over 80% sensitivity.

**Figure 2.**
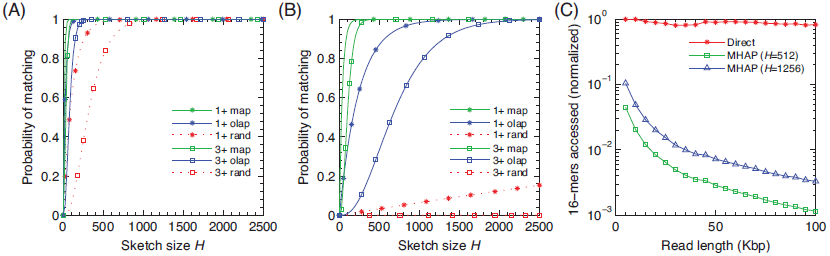
Simulated MHAP performance for various sketch sizes and read lengths. Reads were randomly extracted from the human reference genome and errors were introduced to simulate a SMRT sequencing error model (11.88% insertion, 1.83% deletion, and 1.29% substitution)^17^. (A) Probability of detecting ≥1 or ≥3 matching min-mers for *k*=10 and various sketch sizes. (B) Probability of detection for *k*=16. Match types are divided into: unrelated sequences (rand), overlapping reads (olap), and reads mapped to a perfect reference (map). The expected Jaccard similarity between a pair of random and non-random reads was estimated based on 50,000 independent trials and equation (9) in the Methods. (C) The total number of 16-mers processed for a direct lookup approach versus MHAP using sketch sizes of 512 and 1256 across varying read lengths. Plot is normalized by the maximum value observed during the simulations.

The efficiency of MHAP improves with increased read length. Figure 2c compares the total number of *k*-mers counted during MHAP overlapping to a direct approach that exactly measures the Jaccard similarity between two reads without using sketches (Methods). For a fixed number of total bases sequenced, and a minimum 20% overlap length, increasing read length has no effect on the direct approach since it must consider all *k*-mers. However, the number of min-mer comparisons performed by MHAP decays rapidly with increasing read length, since the complexity is governed by the sketch size (a constant) and the number of reads (which decreases for increasing read length, Supplementary Table S1). Thus, the efficiency of MHAP is expected to improve with the increasing read length and accuracy of future long-read sequencing technologies.

### MHAP overlapping performance

In addition to speed, MHAP is a highly sensitive overlapper. We evaluated the sensitivity and specificity of MHAP versus BLASR^32^, the only other aligner currently capable of overlapping SMRT reads. BWA-MEM^33^, SNAP^34^, and RazerS^35^ were also evaluated, but their current versions were unable to reliably detect noisy overlaps (Supplementary Note 2). The performance of MHAP and BLASR was evaluated by comparing detected overlaps to true overlaps inferred from mapping reads to their reference genomes (Methods). First, the methods were evaluated across multiple parameter settings and sequencing chemistries (Supplementary Figs. S1–S2, Table S2). Importantly, MHAP sensitivity is tunable based on the size of *k*, the sketch size, and the Jaccard similarity threshold. Based on these results, two MHAP parameter sets were chosen that balanced speed with accuracy, and the comparisons were repeated on all datasets (Table 1). To achieve adequate sensitivity on large, repetitive genomes, BLASR required parameter settings that drastically increased runtime. For efficiency, BLASR only considers a subset of alignments (controlled by the *bestn* and *nCandidates* parameters), resulting in missed overlaps for highly repetitive genomes. In contrast, MHAP considers all alignments and was consistently accurate across all genomes tested, while being orders of magnitude faster than BLASR.

**Table 1.** **Overlapping sensitivity and specificity for MHAP and BLASR** Organism: The genome analyzed. Program: the program being run. Sensitivity: the percentage of true overlaps identified by the program. Specificity: the percentage of true negatives correctly identified by the program. PPV (Positive Predictive Value): the percentage of true overlaps out of all overlaps reported. In all cases, a maximum of 1Gbp was randomly selected and mapped to the reference genome. BLASR was run with default parameters and *bestn* set to sequencing depth of coverage (BLASR 1C), 10 times coverage (BLASR 10C), and 100 times coverage (BLASR 100C). MHAP was run with default parameters (*k=16, min. matches=3, min. Jaccard=0.04*) and sketch sizes of 512 (MHAP 512) and 1256 (MHAP 1256). True positives and false negatives were estimated to ±1% (Methods, Supplementary Note 8). Smith-Waterman alignment was used to confirm overlaps not detected by reference mapping (Supplementary Note 8). For *D. melanogaster*, BLASR with *bestn*=100C was terminated after seven days.

| Organism | Program | Sensitivity | Specificity | PPV | Time (CPU h) | Overlap Mem (GB) |
| --- | --- | --- | --- | --- | --- | --- |
| *E. coli* K12 | BLASR 1C | 89% | 100% | 100% | 21.53 | 3.71 |
|  | BLASR 10C | 95% | 100% | 100% | 86.84 | 3.71 |
|  | BLASR 100C | 95% | 100% | 100% | 706.32 | 4.40 |
|  | MHAP 512 | 65% | 100% | 99% | 2.27 | 20.14 |
|  | MHAP 1256 | 85% | 100% | 99% | 3.61 | 22.34 |
| *S. cerevisiae* W303 | BLASR 1C | 16% | 100% | 100% | 71.43 | 5.42 |
|  | BLASR 10C | 35% | 100% | 100% | 434.30 | 5.71 |
|  | BLASR 100C | 87% | 100% | 99% | 2,832.67 | 13.20 |
|  | MHAP 512 | 70% | 100% | 98% | 18.58 | 8.78 |
|  | MHAP 1256 | 83% | 100% | 98% | 31.44 | 10.40 |
| *A. thaliana* Ler-0 | BLASR 1C | 2% | 100% | 100% | 30.92 | 8.93 |
|  | BLASR 10C | 11% | 100% | 100% | 82.24 | 5.85 |
|  | BLASR 100C | 68% | 100% | 100% | 445.27 | 6.03 |
|  | MHAP 512 | 69% | 100% | 100% | 13.34 | 9.86 |
|  | MHAP 1256 | 88% | 100% | 100% | 21.27 | 11.10 |
| *D. melanogaster* ISO1 | BLASR 1C | 75% | 100% | 100% | 166.81 | 11.70 |
|  | BLASR 10C | 89% | 100% | 100% | 878.15 | 11.44 |
|  | BLASR 100C | - | - | - | - | - |
|  | MHAP 512 | 62% | 100% | 99% | 9.08 | 10.49 |
|  | MHAP 1256 | 83% | 100% | 98% | 15.73 | 11.58 |
| Human CHM1htert | BLASR 1C | 41% | 100% | 99% | 34.20 | 6.60 |
|  | BLASR 10C | 59% | 100% | 96% | 42.50 | 4.68 |
|  | BLASR 100C | 71% | 100% | 93% | 189.45 | 5.02 |
|  | MHAP 512 | 64% | 100% | 93% | 9.37 | 9.20 |
|  | MHAP 1256 | 77% | 100% | 92% | 17.19 | 10.72 |

### SMRT sequencing and assembly

MHAP overlapping enables efficient and complete assembly of large genomes. To demonstrate, we integrated MHAP into the Celera Assembler^36^ PBcR^15, 17^ hierarchical assembly pipeline, and assembled the genomes of *Escherichia coli* K12, *Saccharomyces cerevisiae* W303, *Drosophila melanogaster* ISO1, *Arabidopsis thaliana* Ler-0, and the haploid human cell line CHM1htert^37^ from high-coverage SMRT sequencing data (85X, 117X, 121X, 144X, and 54X coverage, respectively, Supplementary Note 3, Supplementary Table S3)^38^. This new pipeline is referred to as PBcR-MHAP, while the previous pipeline based on BLASR is PBcR-BLASR. Also included in PBcR-MHAP is a new consensus module, FalconSense, for correcting the noisy reads after overlapping (Methods). Table 2 details the relative performance of PBcR-MHAP on all datasets. Assembly continuity is measured using the traditional N50 metric (half the genome size is contained in contigs of length *N* or greater) and an “assembly performance” metric defined by Lee *et al.* as the N50 contig length divided by the N50 length of the reference segments (observed N50 vs. idealized N50)^12^. In all cases compared, MHAP overlapping produced comparable or improved assembly results in less time that previous approaches. As expected, speedups were most pronounced for larger genomes and read lengths, with a ∼600-fold speedup observed for *D. melanogaster.*

**Table 2.**
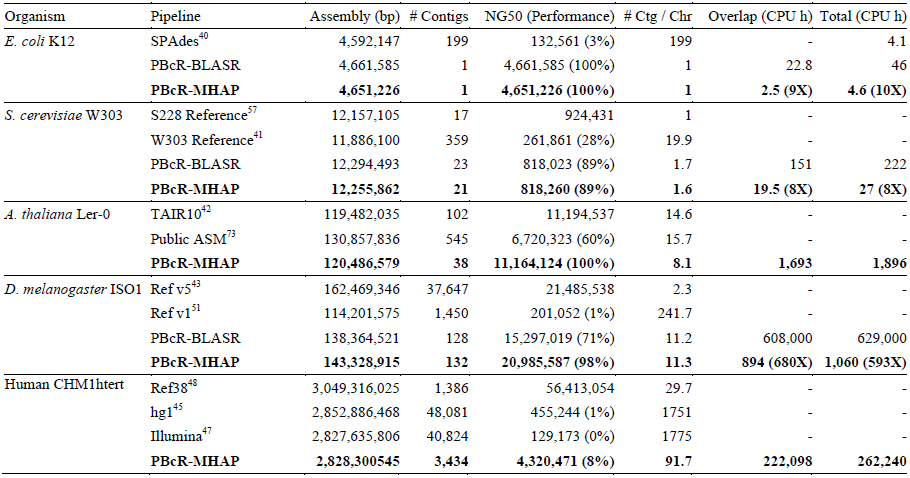
**Continuity and performance of long read assembly with MHAP** Organism: The genome being assembled. Pipeline: pipeline used for correction and assembly. Assembly: the total number of base pairs in all contigs (only contigs containing at least 50 reads are included in all PBcR results). # Contigs: The number of contigs >200 bp. NG50: *N* such that 50% of the genome is contained in contigs of length ≥*N* where the genome size is set to reference length, excluding unknown sequences. Assembly performance is the contig N50 divided by the N50 of the reference segments12. The genome sizes were estimated from the reference to be 4,639,675 for *E. coli* K12; 12,157,105 for *S. cerevisiae* W303; 119,482,035 for *A. thaliana* Ler-0; 129,663,327 for *D. melanogaster* ISO1; and 3,101,804,741 for human*. Average # Contigs / Chromosome:* Excluding unassigned scaffolds, average number of contigs >200 bp per chromosome. Average for assemblies are based on alignments to the reference genome, while averages for reference genomes (and N50) are based on splitting at three or more consecutive Ns. Overlap Time: CPU hours to compute overlaps using BLASR or MHAP (with fold speedup versus BLASR, where applicable). All timing was performed on AMD 2.4GHz 6136 processors. Total time (CPU h): total time taken to assemble the genome. All overlapping steps were constrained to 32GB of RAM per computational node.

Using PBcR-MHAP, microbial genomes can be completely assembled from long reads in roughly the same time required to generate incomplete assemblies from short reads. For example, PBcR-MHAP was able to accurately resolve the entire genome of *E. coli* K12 using 85X of SMRT reads in 4.6 CPU hours, or 20 minutes using a modern 16-core desktop computer. In comparison, the state-of-the-art^39^ SPAdes assembler^40^ required 4.1 CPU hours to assemble 85X Illumina reads from the same genome. Both short- and long-read assemblies are highly accurate at the nucleotide level (>99.999%), but the short-read assembly is heavily fragmented and contains more structural errors (Supplementary Table S4, Supplementary Fig. S3). Our initial SMRT assembly does contain more single-base insertion/deletion (Indel) errors, but polishing it with Quiver^19^ (requiring an additional 6.6 CPU hours) resulted in the lowest number of consensus errors of all assemblies (11 vs. 96 for SPAdes).

*S. cerevisiae* W303, the smallest eukaryotic genome assembled here, was sequenced using the P4C2 chemistry, which produces shorter reads than P5C3. Nonetheless, the resulting PBcR-MHAP assembly produced fewer contigs than PBcR-BLASR, ran eight times faster, and required less than two hours on a desktop computer. Despite being from a different strain, the assembly is largely syntenic with the *S. cerevisiae* S228 reference (Supplementary Fig. S4). Compared to a previously published reference-guided assembly of this strain^41^, the PBcR-MHAP *de novo* assembly achieves a 4-fold improvement in N50 and assembles 12 out of 16 chromosomes without gaps. Thus, even using the older P4C2 chemistry, our assembly approaches perfect continuity and represents the best assembly of *Saccharomyces cerevisiae* W303 to date, including both hybrid and reference-assisted assemblies^12, 41^.

As predicted, greater performance gains were observed for the larger, more complex genomes of *A. thaliana* Ler-0 and *D. melanogaster* ISO1 sequenced using the longer P5C3 chemistry (Table 2). Compared to the *A. thaliana* Col-0 version 10 reference^42^, which has undergone continued improvement since its initial sequencing in 2000, our assembly of Ler-0 is more continuous and contains an average of 5 fewer gaps per chromosome. The Ler-0 assembly structurally agrees with the Col-0 reference, with the exception of a few strain-specific variations (Supplementary Fig. S5). Similarly, the *D. melanogaster* ISO1 assembly achieves startling continuity and widespread agreement with the version 5 reference^43^ (Supplementary Note 4, Supplementary Fig. S6). For example, chromosome arm 3L is fully spanned by a single 25Mbp contig (Fig. 3), and on average our assembly is significantly more contiguous than the original Sanger-based assembly of *D. melanogaster*^36^, averaging just 5 contigs per autosomal chromosome arm (2L, 2R, 3L, 3R, 4). As a result of this outstanding continuity, our assembly potentially resolves 52 of 124 (42%) gaps in the version 5 reference that have persisted for over a decade of finishing (Supplementary Note 4, Supplementary Table S5). Half of these putative gap closures match the estimated gap size in the reference, but require further validation to confirm they represent true gap closures.

**Figure 3.**
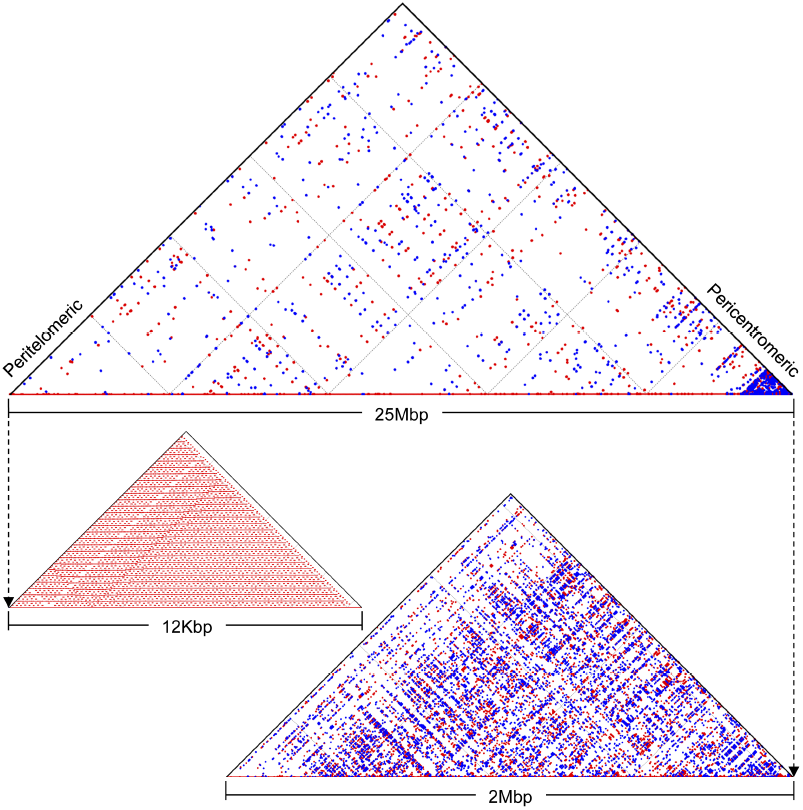
**Single-contig assembly of *D. melanogaster* chromosome arm 3L** A singe ∼25Mbp contig from the PBcR-MHAP *D. melanogaster* assembly covers the full euchromatic region of chromosome arm 3L. (Top) All 100bp exact repeats across the length of the 3L assembly are shown using a self-alignment dotplot. Red dots indicate forward repeats, and blue dots inverted repeats. Points nearer to the hypotenuse indicate repeat copies nearer to each other in the genome. (Bottom left) All 20bp exact repeats are shown for the first 12Kbp of the assembly illustrating a peritelomeric tandem repeat. (Bottom right) All 20bp exact repeats are shown for the last 2Mbp of the assembly, which is comprised of an elevated repeat density, characteristic of the pericentromeric region.

Despite the original Drosophila Genome Project's 2Kbp, 10Kbp, and BAC inserts, the *contig* N50 of the SMRT assembly is greater than the *scaffold* N50 of the Sanger assembly (21Mbp vs. 14Mbp). This result demonstrates that sufficient coverage of long reads can independently resolve repeats in eukaryotic genomes, eliminating the need for read pairs. Lastly, only a few days was required to assemble the genomes of *A. thaliana* and *D. melanogaster* using PBcR-MHAP on a single desktop computer, demonstrating that reference-grade assembly of 100Mbp eukaryotes is now possible without requiring large computing clusters.

### A *de novo* human assembly using long reads

The human genome has long been regarded as the pinnacle of whole-genome shotgun sequence assembly^44^, and is clearly the most widely studied genome with enormous resources dedicated to sequencing, assembling, and finishing^45–47^. As a final test of PBcR-MHAP, we assembled 54X SMRT sequencing reads from the haploid human cell line CHM1htert^37^. We compared our assembly to the human GRCh38 reference^48^, as well as to a reference-guided assembly of CHM1 that utilized BAC-tiling and Illumina sequencing^47^ (Supplementary Note 5, Supplementary Fig. S7).

The contig N50 of our long-read *de novo* assembly of CHM1 is an order of magnitude larger than both the CHM1 Illumina assembly and the original Sanger-based assemblies of human (Fig. 4, Table 2, Supplementary Figs. S8-S10). Based on a comparison to the GRCh38 reference, the average number of contigs per chromosome in our assembly is 92. Not only does our assembly exceed all previous *de novo* assemblies of human, but it also has the potential to improve the current human reference. Our assembly potentially resolves 51 of 819 (6%) annotated gaps in GRCh38 (Fig. 4, Supplementary Note 4, Supplementary Table S5). Of these putative gap closures, 16 match the estimated gap size in the reference, but require further validation to confirm they represent true gap closures.

**Figure 4.**
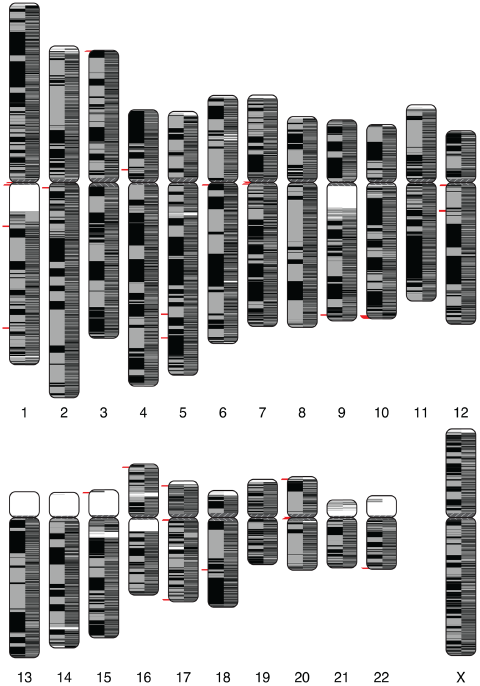
**Continuity and putative GRCh38 gap closures of the human CHM1htert assembly** Human chromosomes are painted with assembled CHM1 contigs (using the coloredChromosomes package^72^). Alternating shades indicate adjacent contigs, so each vertical transition from gray to black represents a contig boundary or alignment breakpoint. The left half of each chromosome shows the PBcR-MHAP assembly of the SMRT dataset and the right half shows the Illumina-based assembly^47^. The SMRT assembly is significantly more continuous, with an average of less than 100 contigs per chromosome. Putative gap closures by the SMRT assembly are shown as red dashes next to the spanned gap position. Most fall near the telomeres and centromeres. Tiling gaps, in white, typically coincide with regions of uncharacterized reference sequence (i.e. long stretches of N's) for which no contigs could be mapped.

One example of a historically difficult region to assemble in the human genome is the major histocompatibility complex (MHC), which plays an important role in immunity^49^. In contrast to the CHM1 Illumina assembly, which breaks the MHC region into over 60 contigs, PBcR-MHAP assembles 97% of this region in just 2 contigs. Compared to the Illumina assembly, a total of 21 (5%) additional MHC genes are correctly reconstructed into a single piece by our assembly.

### Assembly validation and repeat resolution

To validate the resulting assemblies, we compared each assembly to the closest available reference genome using dnadiff^50^ (Supplementary Note 4, Supplementary Table S4). All assemblies are structurally concordant with the reference sequences (Supplementary Figs. S3–S7). However, only the *E. coli* and *D. melanogaster* datasets were generated from the same strain as the reference, and detailed validation was limited to these genomes. The *D. melanogaster* SMRT reads were sequenced from the same subline of ISO1 used by the Drosophila Genome Project since 2000^51^ providing the opportunity to validate SMRT sequencing of a eukaryotic genome.

Supplementary Table S4 provides GAGE^52^ accuracy metrics for the *E. coli* and *D. melanogaster* assemblies. Averaging across these two assemblies, and ignoring the effects of heterzyogsity, we estimate PBcR-MHAP consensus accuracy to be 99.9%, corresponding to a Phred Quality Value (QV) of 30. Further polishing the raw assemblies with Quiver increased the average base accuracy to 99.99% (QV40), but at the expense of added runtime. For example, Quiver polishing of the 143Mbp *D. melanogaster* assembly required a total of 498 CPU hours. Thus, true diploid assembly and consensus polishing remains an area for improvement. Alternatively, short-read resequencing could be used to polish long-read draft assemblies and call heterozygous alleles via mapping.

We further analyzed the completeness of the *D. melanogaster* chromosomes and gene sequences^53^ in our assembly (Supplementary Note 6). A total of 2,884 high-quality Nucmer^54^ alignments cover 122.9Mbp (95.7%) of the 129.7Mbp version 5 reference genome^43^. Of these alignments, 114.8Mbp cover known euchromatic sequence and 8.1Mbp cover known heterochromatic sequence. We further mapped 17,294 annotated genes (full-length, including introns) to our assembly, identifying a total of 16,604 (96%) genes contained in a single alignment to a single contig. Of these, 15,780 were reconstructed at over 99% identity and 7,299 were reconstructed with perfect identity. After Quiver, these numbers increased to 16,776 genes in a single alignment (97%), 16,751 over 99% identity, and 14,824 perfectly reconstructed.

Because repeats represent the greatest challenge to assembly, we also analyzed the completeness of *D. melangaster* transposable element (TE) families. TEs in *D. melanogaster* represent a large fraction of annotated repeats and vary over a wide range of sizes and levels of sequence diversity^55^. To assess TE resolution in our assembly, we replicated the analysis of a recent study that assembled *D. melanogaster* from Illumina Synthetic Long Reads (Moleculo) using the Celera Assembler^56^. Of 5,425 annotated TE elements in the euchromatic arms^55^, 5,201 (96%) are contained in a single contig by our assembly and the majority aligned perfectly to the reference (4,026 pre-quiver; 4,984 post-quiver). Our assembly also resolves 96% (132 of 138) of copies from the highly-abundant *roo* TE family (37 at 100% identity; 93 post-Quiver), in contrast to the results of McCoy *et al.*^56^, where only 5.2% of *roo* copies were resolved. For the *juan* family, with less than 0.01% divergence between copies, a total of 7 of 11 copies were reconstructed at perfect identity (11 of 11 post-Quiver). We conclude that the SMRT assembly accurately reconstructs a significantly higher fraction of complex TEs than Illumina Synthetic Long Reads. In addition, the high error rate of SMRT sequencing does not appear to prohibit the accurate reconstruction of TE sequences, even from highly abundant TE families with many identical copies interspersed throughout the genome.

### Improved telomere assemblies

Because SMRT sequencing generates long reads without the need for cloning or amplification, it is possible to better reconstruct the repeat-rich heterochromatic regions of eukaryotic chromosomes. This is a distinct advantage compared to previous sequencing methods, for which heterochromatin sequencing was thought to be impossible because of cloning biases or short read lengths. As proof of principle, we evaluated the ability of long-read sequencing to reconstruct heterochromatic sequences in the telomeric regions of *S. cerevisiae*^57^ and *D. melanogaster*^43^ (Supplementary Note 7). Telomeres play important roles in chromosome replication in all eukaryotic genomes, and in humans their loss has been associated with disease^58^, but these sequences are typically missing from *de novo* assemblies in eukaryotes. For example, it was not until six years after the initial shotgun assembly that the *D. melanogaster* reference genome began to include telomeric sequence^59^.

As in other genomes, *S. cerevisiae* telomeres serve to protect the chromosome and aid in chromosome repair. Additionally, they affect the transcription of nearby genes^60^. Using the annotations in the reference genome, we mapped the telomeric repeats to our assembly of *S. cerevisiae* W303 (Supplementary Note 7). We identified 9 chromosomes where a single contig included an alignment comprising at least 50% of the terminal telomeric repeat on both the left and right ends, indicating that a majority of the 16 chromosomes were completely resolved from telomere to telomere. In the remaining cases, 4 chromosomes were composed of more than one contig containing the telomeres, and 2 chromosomes did not extend into the telomeres (or the telomeric sequence did not match the reference). This is a significant improvement over the current *S. cerevisiase* W303 genome^41^, where only one chromosome is spanned from end to end and only 5 chromosome ends have been annotated.

In contrast to the simple telomeric repeats of *S. cerevisiae* and other eukaryotes, *D. melanogaster* telomeres are composed of head-to-tail arrays of three specialized retrotransposable elements (*Het-A*, *TART*, and *Tahre*) and clusters of telomere-associated sequences (TASs)^59, 61, 62^. Because telomeric TEs preferentially transpose to chromosome ends, they are virtually absent from the euchromatic regions^59^. By mapping repeat families from RepBase^63^ and a recent *D. melanogaster* repeat study^64^, we identified repeat arrays characteristic of *D. melanogaster* telomeres (Supplementary Note 7). A total of 24 telomeric contigs are present in our assembly. One contig, corresponding to chromosome 2R, contains both subtelomeric (*HetRp_DM*) and telomeric sequence, fully capturing the transition from euchromatin to heterochromatin. This contig extends 80Kbp beyond the end of the reference assembly, and represents the first time the full telomeric transition sequence has been identified for this chromosome. Chromosome arms 2L, 3R, and X also have large contigs containing telomeric repeats extending past the end of the current reference sequence, indicating additional regions of the reference that could be improved by our *de novo* assembly.

### Cloud computing for large-genome assembly

Exponentially lower costs have democratized DNA sequencing, but assembling a large genome still requires substantial computing resources. Cloud computing services offer an alternative for researchers that lack access to institutional computing resources. However, the cost of assembling long-read data using cloud computing has been prohibitive. For example, using Amazon Web Services (AWS), the estimated cost to generate the *D. melanogaster* PBcR-BLASR assembly is over $100,000 at current rates, an order of magnitude higher than the sequencing cost. With MHAP, this cost is drastically reduced to under $300. To expand access to the PBcR-MHAP assembly pipeline, we have provided a free public AWS image as well as supporting documentation for non-expert users that reproduces the *D. melanogaster* assembly presented here in less than 10 hours using AWS. Allocating additional compute nodes, which would marginally increase costs, could further reduce assembly time. For *E. coli,* the total cost of PBcR-MHAP assembly and Quiver polishing is currently less than $2. With MHAP, assembly costs are now a small fraction of the sequencing cost for most genomes, making long-read sequencing and assembly more widely accessible.

## Discussion

We have demonstrated that it is possible to assemble large genomes from noisy, long reads. Like was previously shown for microbial genomes, assembly of eukaryotic genomes using SMRT sequencing can automatically produce reference-grade genomes. In the best cases, entire chromosome arms assemble into single-piece assemblies from telomere to centromere. For the few remaining gaps, long-read assemblies could be paired with super-long linking information as generated by optical^65, 66^ or chromatin interaction maps^67–69^. These complementary scaffolding approaches could be used to span centromeres, resolve entire chromosomes, and phase haplotypes to generate truly complete assemblies.

Our results indicate that probabilistic alignment methods are well suited to address the read length and error rate of single-molecule sequencing. A number of strategies have been previously developed to find similarities in high dimensional data, and chief among them are probabilistic dimensionality reduction approaches^23, 70, 71^. Such algorithms trade the guaranteed accuracy of a deterministic method for a much faster solution with bounded error. By producing high-quality assemblies in a fraction of the time, MHAP demonstrates that this tradeoff is acceptable for the overlapping problem, where sequencing redundancy compensates for loss in alignment accuracy. Further, the long reads produced by single-molecule sequencing allow for a coarser estimate of similarity than traditional dynamic programming or direct *k*-mer matching.

MHAP can serve as drop-in replacement for current overlapping methods. The sensitivity of MHAP is well suited for overlapping diploid or polyploid genomes; however, the genomes presented here are either inbred or haploid and current assemblers struggle to reconstruct genomes with structurally divergent alleles. In addition to SMRT sequencing, MHAP is likely to be suitable for nanopore sequencing^10^, which is expected to have similar read length and error characteristics. As a strategy, MinHash sketches are also applicable to reference alignment, sequence clustering, and alignment-free distance estimation. Future work includes evaluating the applicability of MinHash to these other areas. Fast and sensitive methods are needed not just for long-read overlapping, but to address the ever-expanding scale of genomic data for all applications.

In addition to demonstrating the potential of probabilistic alignment and long-read sequencing, we have freely provided high-quality assemblies of human CHM1 and the important model organisms *S. cerevisiae* W303, *A. thaliana* Ler-0, and *D. melanogaster* ISO1. We hope these assemblies will assist in the continued finishing of these important reference genomes.

## Acknowledgements

We are indebted to Casey Bergman of the University of Manchester for his considered advice throughout this project and editing of an early version of this manuscript. We also thank Pacific Biosciences and all those involved in generating and freely releasing the data analyzed here. The contributions of SK and AMP were funded under Agreement No. HSHQDC-07-C-00020 awarded by the Department of Homeland Security Science and Technology Directorate (DHS/S&T) for the management and operation of the National Biodefense Analysis and Countermeasures Center (NBACC), a Federally Funded Research and Development Center. The views and conclusions contained in this document are those of the authors and should not be interpreted as necessarily representing the official policies, either expressed or implied, of the U.S. Department of Homeland Security. In no event shall the DHS, NBACC, or Battelle National Biodefense Institute (BNBI) have any responsibility or liability for any use, misuse, inability to use, or reliance upon the information contained herein. The Department of Homeland Security does not endorse any products or commercial services mentioned in this publication.

## Author Contributions

KB and SK conceived, designed, and implemented the MHAP algorithm. CSC and JD conceived, designed, and implemented the consensus algorithms. SK ran and analyzed the genome assemblies. JML coordinated data release and assisted with pipeline executions. CSC and SK performed cloud-computing experiments. KB, SK, and AMP drafted the manuscript. AMP coordinated the project. All authors read and approved the final manuscript.

## Competing Interests

CSC, JD, and JML are stockholders and employees of Pacific Biosciences. KB, SK, and AMP have no competing interests.

**Correspondence** All correspondence and requests for materials should be addressed to Sergey Koren.

**Availability** The latest version of Celera Assembler (including PBcR) is available from http://wgs-assembler.sourceforge.net. The software and data used for this manuscript (including the AWS recipe) are available from http://www.cbcb.umd.edu/software/PBcR/MHAP. Assemblies are available at the above website and will be submitted to GenBank.

## Methods

In order to provide context for the MHAP algorithm, we first describe the evolution of read overlapping approaches, and then describe how MHAP can be viewed as another evolution of existing techniques. We start by demonstrating how long reads can degrade the performance of current approaches. Then, we introduce the notion of Jaccard similarity and describe how it depends on read length and error rate. Next, we show how Jaccard distance can be efficiently computed and how it is implicitly measured by current approaches. Finally, we introduce the concept of MinHash and demonstrate how to apply it to noisy, long reads.

### Background

An overlap is typically defined as maximally scoring alignment between two strings that allows arbitrary orientation and offset of the reads^74^. For two reads *S_1_* and *S_2_*, both of length *O(L)*, the fastest method for determining their optimal alignment is Smith-Waterman (SW) dynamic programming, which has a computational complexity of O(*L*^2^)^75^. Thus, to naively find all overlapping pairs of *N* reads would take O(*N^2^L^2^*).

In order to reduce the number of pairs that need to be directly evaluated by SW, each read can be broken into a set of *k*-length substrings (*k-*mers) by sliding a *k*-sized window along the length of the read. Thus, a read *S* of length *L* is represented by a set of *K*=*L*-*k*+1 *k*-mers. The *k*-mers of *S* can be indexed in a hash table (or suffixes of *S* in a suffix array) in O(*NK*) time^76^. This index can be used to filter out potentially non-matching pairs faster than a direct O(*N^2^*) comparison. To compare two reads, the number of shared *k*-mers is computed by the counting how many *k*-mers in a query read match *k*-mers in other reads^32, 35, 36, 77^. The overall time, when comparing against multiple reads, can be improved by using a suffix array to index all existing reads^32, 54^. Overlapping algorithms designed for high-identity reads, such as the original Celera Assembler overlapper^36^, may trigger a banded SW search based on a single shared *k*-mer. However, using any of these approaches, the complexity is proportional to the read length.

For reasons that will become apparent below, we focus our discussion on the hash table indexing approach, though similar reasoning applies to the suffix array approaches. In the case of hash table indexing, all *k*-mers are first hashed into an integer fingerprint for easier indexing and faster comparison. We define this set of *K*=*L*-*k*+1 integers as *Γ*(*S*), where |*Γ*(*S*)|=*K* is the set's cardinality. Here we assume that the integer size is large enough that a chance of a random collision is negligible. The integers from all strings are hashed into a table such that each hash table bucket will contain a list of all string indices that contain that specific integer fingerprint. In order to find all the strings that have at least *w* shared *k*-mers with *S*, each element in *Γ*(*S*) is found in the hash table, and a count table is maintained for any read that has at least one matching *k*-mer. Only the string in the count table that have ≥*w* counts are returned. Observe that the time to maintain the count table is directly related to the sum of all counts in the count table and is an additional cost to the hash table lookup of all *k*-mers.

We refer to the above as a “direct” method and demonstrate how using noisy, long reads can rapidly degrade its performance. While straightforward, this approach is actively used in efficient alignment algorithms for short-read sequencing^34^ and is a viable alternative to compressed index alternatives like suffix arrays^78^. As an alternative, we present a probabilistic dimensionality reduction and filtering algorithm that theoretically improves runtime and storage requirement over direct methods or suffix-array approaches such as BLASR^32^.

### Matching *k*-mer probabilities

Given two random reads generated from a finite alphabet set Σ (e.g. {A,C,G,T}) of length *L* and an integer *k*≤*L,* each read contains *K k*-mer fingerprints, where the expected number of fingerprints that are shared by these two reads, *E*[*X_r_*], is equal to the probability that a random *k*-mer is in the first read,

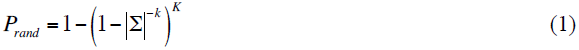

multiplied by the number of *k*-mers in the second read,

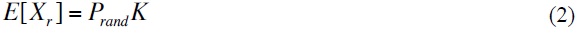

Similarly, the expected number of *k-*mers that are shared by two reads of arbitrary length within an overlapping *M*≤*K* size *k*-mer region, *E*[*X_c_*], corrupted by *ε* error at each position, is related to the probability that an overlapping *k*-mer is not corrupted in both reads,

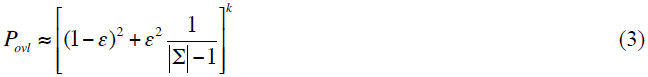

(assuming substitution errors, for insertions or deletions the probability is slightly different, but on the same order), multiplied by the length of the overlap

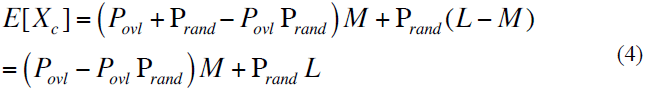

The sensitivity and accuracy of using *k*-mer counts for filtering overlapping pairs, using any data structure, is dependent on how accurately we can classify if a *k-*mer count between any two reads is coming from either *X_r_* or *X_c_*.

Given the small DNA alphabet, from eq. (2) and (4), we observe that: (i) for large *L* the value of *E*[*X_r_*] starts to approach that of *E*[*X_c_*], and determining if two reads are actually overlapping (or random matches) based on the *k*-mer count becomes more error-prone; (ii) the number of *k*-mer hash lookups required for each read grows with *L*; (iii) the percentage of actual reads that collide for each specific *k*-mer lookup grows as 1− (1− 4^−^*^k^*)*^L^*. So, for example, while for *K*=200 one expects on average that approximately 0.02% of total sequences match any given 10-mer, for *K*=50,000 this grows to approximately 5% of total reads, with the probability of having a 10-mer match for every read at least once, when performing *K* 10-mer lookups, ≈ 100%. Unless *k* is large enough for a specific *L*, the computational complexity of looking up a length *L* read in a data structure is not *O(K)*, but rather O(*KN)*. In other words, the computational cost is related to the time it takes to maintain the count table, rather than the overhead of lookups in the data structure storing the *k*-mers. Therefore, the resulting number of operations required for an all-to-all lookup is *O(KN^2^),* with the constant decreasing exponentially with *k*. Importantly, the maximum *k* is limited by the error rate of the reads and the size of the overlap. Thus, *ε*, *L,* and *M* bound the performance of an all-to-all lookup, regardless of the data structure used. Below we will demonstrate that when using MinHash sketches, instead of a full *k*-mer set representation of *S*, we significantly decrease *K*, and by extension the complexity of the computation (Fig. 2a).

### Jaccard similarity

Jaccard similarity^30, 79^ is a measurement related to the actual *k*-mer count, defined as

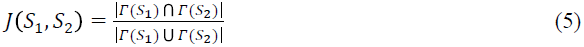

Assuming that the number of repeating *k*-mers (or hash value collisions) is negligible, so both reads have *K k*-mers,

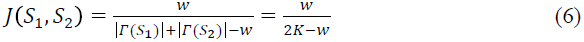

where *w* is the number of *k*-mer matches between *S_1_* and *S_2_*. Assuming *k* is large enough and *ε* is small enough so that the probability of a random match is sufficiently small, the Jaccard similarity between two overlapping reads is proportional to the percentage of sequence length that is shared between the two reads, and thus independent of *L*. Asymptotically, for highly dissimilar matches, the Jaccard similarity approaches *w*/(2*K*), and for highly similar matches it approaches 1.

The Jaccard similarity between two *L*-sized reads, as well as *k*-mer collision count, can be directly computed in O(*L* log *L*) time by using a “sort-merge” algorithm, where we first sort *Γ*(S) of the two reads, and then count the number of matching *k*-mers by performing a merge operation. For multiple comparisons the cost of sorting is amortized, so a direct read comparison complexity drops to O(*L*) time.

### Modified sort-merge algorithm

During read overlapping we are looking for similarity in the overlapping region, rather than the averaged similarity over the full length of the reads, as would be measured by the Jaccard similarity. In addition, assembly algorithms require the approximate position and size of the overlap region for building the read layout.

In order to compute the overlap score and region, we modify the basic sort-merge algorithm by also storing the indices of the *k*-mer position along with the *k*-mer fingerprint. In case a *k-*mer in *S_1_* matches multiple *k-*mers in *S_2_*, we store only the match with the earliest position in the read, thus guaranteeing O(*L*) operations. The position and size of the *S_1_* and *S_2_* overlap is computed using the median difference in position of matching *k*-mers. The *k*-mer counting algorithm is then run again, but now constrained only to the overlapping region, in order to get the final *k*-mer count. Based on the computed *k*-mer count *w* and overlap region size *M*, we define the overlap similarity as

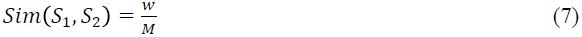

This approach reduces the complexity of computing the similarity between two reads from O(*L^2^*) to O(*L*). Additionally, the O(*L*) runtime is asymptotically faster than the downstream steps, so filtering false matches early does not asymptotically increase the computational complexity of assembly.

### MinHash

Even with the above improvements for computing sequence similarity, we are still left with the problem of computing it for all possible pairs. It is possible to use a hash table or suffix array to accelerate the algorithm^32, 54^. However, we are still faced with a rapid decay in computational efficiency for large *L* (see above). To significantly decrease the number of table lookups per read, as well as the time it takes to build the lookup table, MHAP uses a probabilistic dimensionality reduction approach called MinHash^23^. The efficiency of MinHash versus a direct computation of Jaccard similarity, comes from reducing a read from *K* integer fingerprints, to some smaller, random, possibly repeating vector of *H* fingerprints, where *H*<<*K*. We refer to this compressed representation of the string as a sketch. Since we only consider *H* values for any given read, the storage and read lookup cost decrease proportionally with the size of the sketch (Fig. 2c).

Observe that the probability of *Γ*(*S*_1_) and *Γ*(*S*_2_) having the same minimum value (or *min-mer*) is equal to the probability that a *k*-mer in at least one of the strings also exists in both strings. This is equal to the Jaccard similarity J(*S_1_*,*S_2_*)^23^:

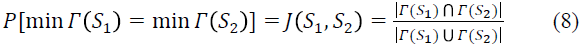

Therefore, given *H* differently seeded hash functions, the probability that the number of min-mers shared by *S_1_* and *S_2_* is at least *x*, can be computed using the cumulative density function of the binomial distribution,

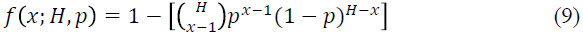

where *p* is the Jaccard similarity of the two sequences (eq. 8). Plots for the percentage of read matches that share at least *x* min-mers, as a function of number of hashes *H,* are shown in Figure 2. The *H* min-mers of *S* forms the sketch of *S*.

### MHAP implementation details

The MHAP algorithm is a two-stage filter that combines the ideas described above. First it filters reads based on shared min-mer counts using MinHash. Then, for all reads that pass the MinHash filter, it again filters matches based on the *k*-mer count for the overlapping region using the sort-merge algorithm to obtain a more accurate similarity estimate. For accuracy, the value of *k* for the second filter can be smaller than the value for the MinHash filter, since we only perform this operation on read pairs that passed the MinHash filter. We refer to the first filter *k*-mer size as *k*_1_, and the second as *k*_2_.

The first filter generates *H* fingerprints for all *k*_1_-mers of a read using the MurmurHash3^80^ hash function implementation in the Guava library^81^ (for larger *k* values a rolling hash can be used). Next, we use the MurmurHash3 fingerprint as a seed into a computationally efficient XORShift random number generator^82^ to generate *H* random fingerprints. Highly repetitive *k*-mers, comprising over 0.001% of the total *k*-mer count, are computed beforehand and ignored. The fingerprint needed for the second filter is computed using the same MurmurHash3, and the fingerprints are sorted and stored in a data structure associated with each *S*, along with the *H* min-mers.

The *h*th min-mer for each read is hashed into the *h*th hash table, which is shared between all reads. The hash tables are maintained in memory, unless there are too many reads. In that case, the computation is treated as a parallel all-to-all computation, where only a subset of reads are hashed and compared to all reads streamed directly from the drive. This subset-to-all comparison is repeated for all non-intersecting subsets, either in parallel or serially, depending on the computation resources available.

For each read we find all the reads that are similar by looking up the *H* min-mers in the hash tables. The *h*th min-mer is looked for in the *h*th hash table, and the counts are aggregated into one list. Any read that has at least *x* min-mers in common with the lookup read is propagated to the second stage filter. The second stage filter is implemented as described above, but to account for the high Indel rate exhibited by SMRT sequencing, the overlap region is extended by 30% on either side.

Since MHAP was designed for complex genomes with large *N* and many repeats, *k*_1_ is set to 16 to decrease the number of false positives in each hash table bucket. *k*=16 is the largest value that can be effectively hashed into a 32-bit fingerprint while providing good sensitivity (Supplementary Table S2). Much larger *k*-mer values would significantly degrade sensitivity, while slightly larger values would double memory usage (requiring a 64-bit fingerprint), without providing significant improvements^17^. Two sketch sizes where chosen (default=512 or sensitive=1256) based on Supplementary Table S2, to achieve sensitivities required for assembly. For the second stage filter, *k*_2_ was empirically set to 12 to avoid false positive matches in 100,000bp reads. Using these settings for SMRT sequencing reads, approximately 40% of overlaps detected using the first filter also pass the second filter. All MHAP parameters are adjustable, and tuning them for specific SMRT chemistries or different sequencing technologies could further improve performance.

### Evaluating sensitivity and specificity

Sensitivity and specificity were evaluated based on overlaps inferred from reference mapping. An automated script was used to evaluate MHAP or BLASR results. To avoid an all-vs-all comparison of reads, random sampling was used to estimate performance. For sensitivity, a random sequence was selected and all sequences with an overlap above a minimum length threshold (2,000bp) were extracted from the reference matches. Any missing overlap was considered a false negative while an existing overlap was a true positive. To compute positive predictive value (PPV), a random overlap above the minimum length threshold was evaluated by comparing it to the reference mapping. To address repeat-induced overlaps, if no reference mapping was found for an overlap, a full local alignment was performed on the overlapping region the overlap marked true if the resulting alignment identity was at least 70% and above the minimum length threshold. The PPV was computed as the number of true overlaps divided by the number of overlaps evaluated. The sample size required to estimate sensitivity, PPV, and specificity to ±1 was calculated using Clopper-Pearson^83^ (Supplementary Note 8).

### Correcting noisy reads

Noisy reads can be accurately corrected using a multiple alignment of reads sampled from the same region of the genome so long as the sequencing errors are random and independent between reads^16, 17^. The correction process consists of two general steps, building a multiple read alignment (MSA) and deciding the correct bases from the MSA. This strategy was first used to correct noisy, long reads using accurate, short reads^17^ and the AMOS make-consensus tool^84^. With the invention of new consensus algorithms, this hierarchical approach has also been successfully applied to the long reads themselves^19^. The PBcR pipeline supports two consensus modules: a slower but more accurate method called PBDAGCon^85^ and a faster but less robust method called FalconSense^86^. PBDAGCon, described previously^19^, uses a directed acyclic graph (DAG) to construct a partial order alignment^87^. Finding an optimized path through the DAG generates a robust consensus. Alternatively, the FalconSense algorithm accelerates consensus generation by never explicitly building a multiple alignment. Instead, FalconSense aligns reads only to a template sequence that is the target of correction^88^. Individual matches and Indels are then tagged and sorted to determine a consensus sequence with high support (Supplementary Note 9, Supplementary Fig. S11)^84, 89^. This read correction algorithm was used for all assemblies presented here, except for human, for which PBDAGCon performed better due to a higher error rate in the raw data.

### Assembling corrected reads

The PBcR pipeline corrects and assembles the input sequences. By default, the longest 40X of data is corrected using all provided sequences and the longest 25X after correction is assembled. These default values were used for all experiments below for both the PBcR-BLASR and PBcR-MHAP pipelines. The spec files and commands used for assembly are reproduced in Supplementary Note 10. Where noted, assemblies were optionally polished using Quiver to optimize base-call accuracy.

### Assembly Validation

All assembled genomes were aligned to their respective reference using Nucmer^54^ and validated using the original GAGE^52^ scripts. To accelerate the alignment of large genomes, Nucmer defaults were modified to increase the minimum seed size and cluster size, as well as to use only unique seeds in the reference genome. Resulting alignments and validation statistics are reported in Supplementary Figures S3–S7 and Supplementary Table S4.

## References

1. Miller, J.R., Koren, S. & Sutton, G. Assembly algorithms for next-generation sequencing data. Genomics 95, 315–327 (2010).

2. Pop, M. Genome assembly reborn: recent computational challenges. Briefings in bioinformatics 10, 354–366 (2009).

3. Nagarajan, N. & Pop, M. Sequence assembly demystified. Nat Rev Genet 14, 157–167 (2013).

4. Bresler, G., Bresler, M. & Tse, D. Optimal assembly for high throughput shotgun sequencing. BMC Bioinformatics 14 **Suppl 5**, S18 (2013).

5. Ukkonen, E. Approximate string-matching with q-grams and maximal matches. Theoretical Computer Science 92, 191–211 (1992).

6. Schatz, M.C., Delcher, A.L. & Salzberg, S.L. Assembly of large genomes using second-generation sequencing. Genome research 20, 1165–1173 (2010).

7. Sanger, F. & Coulson, A.R. A rapid method for determining sequences in DNA by primed synthesis with DNA polymerase. Journal of molecular biology 94, 441–448 (1975).

8. Timp, W. et al. Nanopore Sequencing: Electrical Measurements of the Code of Life. IEEE transactions on nanotechnology 9, 281–294 (2010).

9. Branton, D. et al. The potential and challenges of nanopore sequencing. Nat Biotechnol 26, 1146–1153 (2008).

10. Clarke, J. et al. Continuous base identification for single-molecule nanopore DNA sequencing. Nature nanotechnology 4, 265–270 (2009).

11. Eid, J. et al. Real-time DNA sequencing from single polymerase molecules. Science 323, 133–138 (2009).

12. Lee, H. et al. Error correction and assembly complexity of single molecule sequencing reads. bioRxiv (2014).

13. Loman, N.Q., Josh, Calus, Szymon A P. aeruginosa serotype-defining single read from our first Oxford Nanopore run. http://dx.doi.org/10.6084/m9.figshare.1052996 (2014).

14. Paszkiewicz, K.F., Audrey; Moore, Karen; O'Neill, Paul The second Oxford Nanopore read ever published. figshare. http://dx.doi.org/10.6084/m9.figshare.1060188 (2014).

15. Koren, S. et al. Reducing assembly complexity of microbial genomes with single-molecule sequencing. Genome Biol 14, R101 (2013).

16. Lam, K.K., Tse, D. & Khalak, A. Near-optimal Assembly for Shotgun Sequencing with Noisy Reads. arXiv preprint arXiv:1402.6971 (2014).

17. Koren, S. et al. Hybrid error correction and de novo assembly of single­molecule sequencing reads. Nat Biotechnol 30, 693–700 (2012).

18. Ono, Y., Asai, K. & Hamada, M. PBSIM: PacBio reads simulator-toward accurate genome assembly. Bioinformatics 29, 119–121 (2013).

19. Chin, C.S. et al. Nonhybrid, finished microbial genome assemblies from long-read SMRT sequencing data. Nat Methods 10, 563–569 (2013).

20. English, A.C. et al. Mind the gap: upgrading genomes with Pacific Biosciences RS long-read sequencing technology. PLoS One 7, e47768 (2012).

21. Ribeiro, F.J. et al. Finished bacterial genomes from shotgun sequence data. Genome Res 22, 2270–2277 (2012).

22. Pacific Biosciences. Data Release: Preliminary de novo Haploid and Diploid Assemblies of *Drosophila melanogaster*. http://blog.pacificbiosciences.com/2014/01/data-release-preliminary-de-novo.html (2014).

23. Broder, A.Z. On the resemblance and containment of documents. Compression and Complexity of Sequences 1997. Proceedings, 21–29 (1997).

24. Broder, A.Z. Identifying and filtering near-duplicate documents. Combinatorial pattern matching, 1–10 (2000).

25. Chum, O., Philbin, J. & Zisserman, A. Near Duplicate Image Detection: min-Hash and tf-idf Weighting. BMVC 810, 812–815 (2008).

26. Narayanan, M. & Karp, R.M. Gapped local similarity search with provable guarantees. Algorithms in Bioinformatics, 74–86 (2004).

27. Yang, X. et al. De novo assembly of highly diverse viral populations. BMC Genomics 13, 475 (2012).

28. Rasheed, Z. & Rangwala, H. Mc-minh: Metagenome clustering using minwise based hashing. SIAM International Conference in Data Mining (2013).

29. Roberts, M., Hayes, W., Hunt, B.R., Mount, S.M. & Yorke, J.A. Reducing storage requirements for biological sequence comparison. Bioinformatics 20, 3363–3369 (2004).

30. Jaccard, P. Distribution de la flore alpine dans le Bassin des Dranses et dans quelques régions voisines. Bulletin de la Société Vaudoise des Sciences Naturelles 37, 241–272 (1901).

31. Hamming, R.W. Error detecting and error correcting codes. Bell System technical journal 29, 147–160 (1950).

32. Chaisson, M.J. & Tesler, G. Mapping single molecule sequencing reads using basic local alignment with successive refinement (BLASR): application and theory. BMC Bioinformatics 13, 238 (2012).

33. Li, H. Aligning sequence reads, clone sequences and assembly contigs with BWA-MEM. arXiv preprint arXiv:1303.3997 (2013).

34. Zaharia, M. et al. Faster and more accurate sequence alignment with SNAP. arXiv preprint arXiv:1111.5572 (2011).

35. Weese, D., Holtgrewe, M. & Reinert, K. RazerS 3: faster, fully sensitive read mapping. Bioinformatics 28, 2592–2599 (2012).

36. Myers, E.W. A Whole-Genome Assembly of Drosophila. Science 287, 2196–2204 (2000).

37. Pacific Biosciences. Data Release: ∼54x Long-Read Coverage for PacBio-only De Novo Human Genome Assembly. http://blog.pacificbiosciences.com/2014/02/data-release-54x-long-read-coverage-for.html (2014).

38. Kim, K. et al. Long-read whole-genome shotgun sequence data of five model organisms - E. coli, S. cerevisiae, N. crassa, A. thaliana, and D. melanogaster. In Prep (2014).

39. Magoc, T. et al. GAGE-B: an evaluation of genome assemblers for bacterial organisms. Bioinformatics 29, 1718–1725 (2013).

40. Bankevich, A. et al. SPAdes: a new genome assembly algorithm and its applications to single-cell sequencing. J Comput Biol 19, 455–477 (2012).

41. Ralser, M. et al. The Saccharomyces cerevisiae W303-K6001 cross-platform genome sequence: insights into ancestry and physiology of a laboratory mutt. Open biology 2, 120093 (2012).

42. Arabidopsis Genome Initiative. Analysis of the genome sequence of the flowering plant Arabidopsis thaliana. Nature 408, 796–815 (2000).

43. Hoskins, R.A. et al. Sequence finishing and mapping of Drosophila melanogaster heterochromatin. Science 316, 1625–1628 (2007).

44. Weber, J.L. & Myers, E.W. Human whole-genome shotgun sequencing. Genome Res 7, 401–409 (1997).

45. Lander, E.S. et al. Initial sequencing and analysis of the human genome. Nature 409, 860–921 (2001).

46. Venter, J.C. et al. The Sequence of the Human Genome. Science 291, 1304–1351 (2001).

47. Meltz Steinberg, K., et al. Single haplotype assembly of the human genome from a hydatidiform mole. bioRxiv (2014).

48. Karolchik, D. et al. The UCSC Genome Browser database: 2014 update. Nucleic Acids Res 42, D764–770 (2014).

49. The MHC sequencing consortium. Complete sequence and gene map of a human major histocompatibility complex Nature 401, 921–923 (1999).

50. Phillippy, A.M., Schatz, M.C. & Pop, M. Genome assembly forensics: finding the elusive mis-assembly. Genome biology 9, R55–R55 (2008).

51. Adams, M.D. et al. The Genome Sequence of Drosophila melanogaster. Science 287, 2185–2195 (2000).

52. Salzberg, S.L. et al. GAGE: A critical evaluation of genome assemblies and assembly algorithms. Genome Research 22, 557–567 (2012).

53. St Pierre, S.E., Ponting, L., Stefancsik, R., McQuilton, P. & FlyBase, C. FlyBase 102-advanced approaches to interrogating FlyBase. Nucleic Acids Res 42, D780–788 (2014).

54. Kurtz, S. et al. Versatile and open software for comparing large genomes. Genome biology 5, R12–R12 (2004).

55. Kaminker, J.S. et al. The transposable elements of the Drosophila melanogaster euchromatin: a genomics perspective. Genome Biol 3, RESEARCH0084 (2002).

56. McCoy, R.C. et al. Illumina TruSeq synthetic long-reads empower de novo assembly and resolve complex, highly repetitive transposable elements. bioRxiv (2014).

57. Mewes, H.W. et al. Overview of the yeast genome. Nature 387, 7–65 (1997).

58. Blasco, M.A. Telomeres and human disease: ageing, cancer and beyond. Nat Rev Genet 6, 611–622 (2005).

59. George, J.A., DeBaryshe, P.G., Traverse, K.L., Celniker, S.E. & Pardue, M.L. Genomic organization of the Drosophila telomere retrotransposable elements. Genome Res 16, 1231–1240 (2006).

60. Zakian, V.A. Structure, function, and replication of Saccharomyces cerevisiae telomeres. Annu Rev Genet 30, 141–172 (1996).

61. Biessmann, H. et al. Frequent transpositions of Drosophila melanogaster HeT-A transposable elements to receding chromosome ends. The EMBO journal 11, 4459–4469 (1992).

62. Levis, R.W., Ganesan, R., Houtchens, K., Tolar, L.A. & Sheen, F.-m. Transposons in place of telomeric repeats at a Drosophila telomere. Cell 75, 1083–1093 (1993).

63. Jurka, J. et al. Repbase Update, a database of eukaryotic repetitive elements. Cytogenetic and genome research 110, 462–467 (2005).

64. Koch, P., Platzer, M. & Downie, B.R. RepARK--de novo creation of repeat libraries from whole-genome NGS reads. Nucleic Acids Res 42, e80 (2014).

65. Schwartz, D.C. et al. Ordered restriction maps of Saccharomyces cerevisiae chromosomes constructed by optical mapping. Science 262, 110–114 (1993).

66. Teague, B. et al. High-resolution human genome structure by single-molecule analysis. Proc Natl Acad Sci U S A 107, 10848–10853 (2010).

67. Burton, J.N. et al. Chromosome-scale scaffolding of de novo genome assemblies based on chromatin interactions. Nat Biotechnol 31, 1119–1125 (2013).

68. Kaplan, N. & Dekker, J. High-throughput genome scaffolding from in vivo DNA interaction frequency. Nat Biotechnol 31, 1143–1147 (2013).

69. Selvaraj, S., J, R.D., Bansal, V. & Ren, B. Whole-genome haplotype reconstruction using proximity-ligation and shotgun sequencing. Nat Biotechnol 31, 1111–1118 (2013).

70. Indyk, P. & Motwani, R. Approximate nearest neighbors: towards removing the curse of dimensionality. Proceedings of the thirtieth annual ACM symposium on Theory of computing, 604–613 (1998).

71. Charikar, M.S. Similarity estimation techniques from rounding algorithms. Proceedings of the thiry-fourth annual ACM symposium on Theory of computing, 380–388 (2002).

72. Böhringer, S., Gödde, R., Böhringer, D., Schulte, T. & Epplen, J.T. A software package for drawing ideograms automatically. Online J Bioinformatics 1, 51–61 (2002).

73. PacBio DevNet. Pacific Biosciences DevNet Datasets https://github.com/PacificBiosciences/DevNet/wiki/Datasets (2014).

74. Staden, R. A new computer method for the storage and manipulation of DNA gel reading data. Nucleic Acids Res 8, 3673–3694 (1980).

75. Smith, T.F. & Waterman, M.S. Identification of common molecular subsequences. Journal of molecular biology 147, 195–197 (1981).

76. Altschul, S.F., Gish, W., Miller, W., Myers, E.W. & Lipman, D.J. Basic local alignment search tool. Journal of molecular biology 215, 403–410 (1990).

77. Rasmussen, K.R., Stoye, J. & Myers, E.W. Efficient q-gram filters for finding all epsilon-matches over a given length. J Comput Biol 13, 296–308 (2006).

78. Manber, U. & Myers, G. Suffix arrays: a new method for on-line string searches. 319–327 (1991).

79. Broder, A.Z., Glassman, S.C., Manasse, M.S. & Zweig, G. Syntactic clustering of the web. Computer Networks and ISDN Systems 29, 1157–1166 (1997).

80. Appleby, A. MurmurHash3 http://code.google.com/p/smhasher/wiki/MurmurHash3 (2014).

81. Guava: Google Core Libraries for Java 1.6+. http://code.google.com/p/guava-libraries/ (2014).

82. Marsaglia, G. Xorshift rngs. Journal of Statistical Software 8, 1–6 (2003).

83. Johnson, N.L., Kemp, A.W. & Kotz, S. Univariate discrete distributions, Vol. 444. (John Wiley & Sons, 2005).

84. Pop, M., Phillippy, A., Delcher, A.L. & Salzberg, S.L. Comparative genome assembly. Brief Bioinform 5, 237–248 (2004).

85. Drake, J. & Chin, J. A sequence consensus algorithm implementation based on using directed acyclic graphs to encode multiple sequence alignment. https://github.com/PacificBiosciences/pbdagcon (2014).

86. Chin, J. FALCON: experimental PacBio diploid assembler. https://github.com/PacificBiosciences/falcon/tree/v0.1.3 (2014).

87. Lee, C., Grasso, C. & Sharlow, M.F. Multiple sequence alignment using partial order graphs. Bioinformatics 18, 452–464 (2002).

88. Myers, E.W. AnO (ND) difference algorithm and its variations. Algorithmica 1, 251–266 (1986).

89. Anson, E.L. & Myers, E.W. ReAligner: a program for refining DNA sequence multi-alignments. J Comput Biol 4, 369–383 (1997).

